# Influenza virus ribonucleoprotein complex formation occurs in the nucleolus

**DOI:** 10.1101/2021.02.24.432647

**Authors:** Sho Miyamoto, Masahiro Nakano, Takeshi Morikawa, Ai Hirabayashi, Ryoma Tamura, Yoko Fujita, Nanami Hirose, Yukiko Muramoto, Takeshi Noda

## Abstract

Influenza A virus double-helical ribonucleoprotein complex (vRNP) performs transcription and replication of viral genomic RNA (vRNA). Unlike most RNA viruses, vRNP formation accompanied by vRNA replication is carried out in the nucleus of virus-infected cell. However, the precise subnuclear site remains unknown. Here, we report the subnuclear site of vRNP formation in influenza virus. We found that all vRNP components were colocalized in the nucleolus of virus-infected cells at early stage of infection. Mutational analysis showed that nucleolar localization of viral nucleoprotein, a major vRNP component, is critical for functional double-helical vRNP formation. Furthermore, nucleolar disruption of virus-infected cells inhibited vRNP component assembly into double-helical vRNPs, resulting in decreased vRNA transcription and replication. Collectively, our findings demonstrate that the vRNA replication-coupled vRNP formation occurs in the nucleolus, demonstrating the importance of the nucleolus for influenza virus life cycle.

## Main

Influenza A virus, belonging to the *Orthomyxoviridae* family, possesses eight-segmented, single-stranded, negative-sense RNA as its genome. Each viral genomic RNA (vRNA) segment exists as a ribonucleoprotein complex (vRNP) associated with multiple nucleoproteins (NPs) and a heterotrimeric RNA-dependent RNA polymerase complex composed of PB2, PB1, and PA subunits^1^. The vRNPs, which are flexible double-stranded helices (width, ∼10 nm; length, 30–120 nm)^2^, are responsible for transcription and replication of the vRNAs. On transcription, vRNA is transcribed into 5′-capped and 3′-polyadenylated mRNA by the polymerase complex in a primer-dependent manner. During genome replication, the vRNA is copied into complementary RNA (cRNA) replicative intermediate by *cis*-acting viral polymerase complex, and the cRNA acts as a template for generating more vRNAs, with involvement of a *trans*-activating/trans-acting viral polymerase complex^3,4^. These replication processes are concomitant with ribonucleoprotein complex assembly; the 5′ terminals of the nascent vRNA and cRNA are associated with a newly synthesized viral polymerase complex that is sequentially coated with multiple NPs and assembled into double-helical vRNPs and cRNPs, respectively^5^.

Unlike most RNA viruses, influenza A virus transcribes and replicates its genome in the nucleus of virus-infected cells^6^. Accordingly, influenza A virus transcription, replication, and vRNP formation heavily rely on host nuclear machineries. Upon initiation of vRNA transcription, viral polymerase complex in the vRNP binds to carboxy-terminal domain of host RNA polymerase II (Pol II)^7^. Then, the PB2 subunit binds to 5′-cap structure of host pre-mRNAs or small nuclear/nucleolar RNAs^8,9^, and the PA subunit cleaves and snatches the 5′-capped fragment for use as a primer^10–12^. The requirement of Pol II for initiation of viral mRNA synthesis indicates that the genome transcription takes place in the nucleoplasm, near host Pol II localization. Genome replication and double-helical vRNP formation reportedly involves several intranuclear host factors, such as minichromosome maintenance helicase complex, UAP56, Tat-SF1, and ANP32^13^. Additionally, recent studies have demonstrated the importance of the intranuclear proteins fragile X mental retardation protein (FMRP), protein kinase C, and LYAR in the replication-coupled vRNP assembly^14–16^. However, since these host proteins are localized in different intranuclear domains, subnuclear site of vRNA replication and vRNP formation remains unidentified.

Recently, we showed that a mutant influenza A virus lacking hemagglutinin (HA) vRNA segment efficiently incorporates 18S and 28S ribosomal RNAs (rRNAs) into progeny virions instead of the omitted HA vRNA and that those rRNAs are associated with viral NPs and form vRNP-like structures^17^. Considering that NPs are localized in not only the nucleus but also the nucleolus^18,19^, we hypothesized that assembly of vRNP components into double-helical vRNP occurs in the nucleolus, the site of rRNA transcription, pre-rRNA processing, and ribosomal assembly. Here, we aimed to identify the precise intranuclear site of influenza virus replication-coupled vRNP formation.

## Result

### Nucleolar localization of vRNP components

Given that the nucleolus is a site of vRNP formation, *de novo* synthesized vRNP components (NP, PB2, PB1, PA, and vRNA) should simultaneously exist in the nucleolus of virus-infected cell. To examine their localization in virus-infected cells, Madin–Darby Canine Kidney (MDCK) cells were infected with influenza A virus and fixed over time. Before immunostaining, the virus-infected cells were treated with a protease to remove highly-condensed host nucleolar proteins and RNAs, which is an established method to detect antigens within the nucleolus^20^. Immunostaining with an anti-NP antibody showed that NP was co-localized with nucleolin/C23, a nucleolar marker, 5–7 h post-infections (hpi) (Figure 1a). Localization pattern of the NP at each time point was similar to that without protease treatment except its detection in the nucleolus (Extended Data Figure 1a); newly synthesized NP was detected in the nucleus at 3–5 hpi, the nuclear export was detected at 5–7 hpi, and the majority of NP was detected in the cytoplasm at 7–9 hpi. All viral polymerase subunits, PB2, PB1, and PA, were co-localized with NP in the nucleoli at 5–7 hpi (Figure 1b). Fluorescence *in situ* hybridization (FISH) showed the presence of vRNAs in the nucleolus at 5 hpi (Figure 1c), demonstrating that all RNP components co-exist in the nucleoli at an early stage of infection.

**Figure 1.**
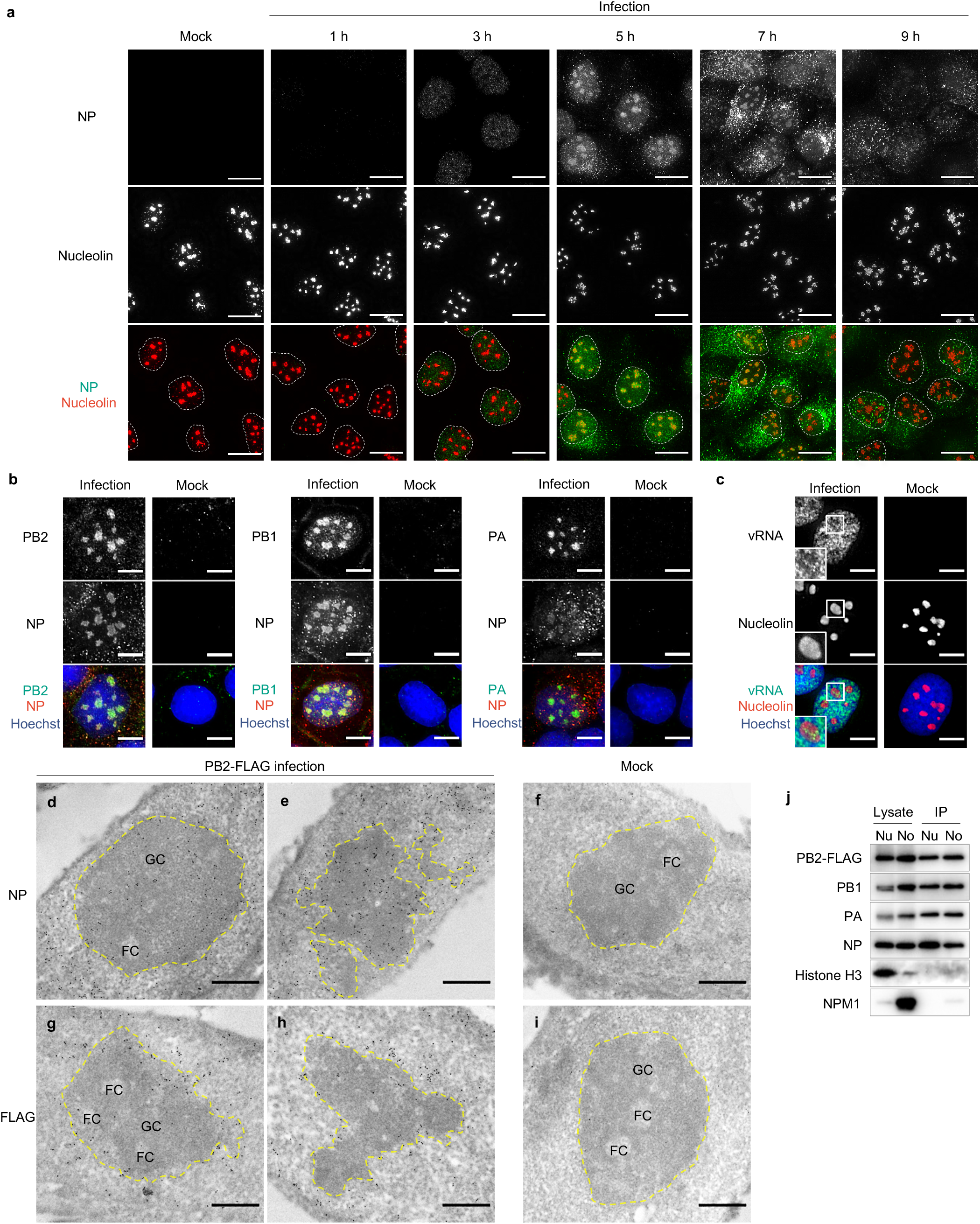
Nucleolar vRNP localization in influenza virus-infected cells. **a,** Subcellular translocation of NPs in mock-infected or influenza virus-infected (MOI=5) cells. NP and nucleolin were immuno-stained after protease treatment of fixed and permeabilized cells. Nuclei are marked by a dashed circle. Scale bars, 20 µm. **b,** Nucleolar co-localization of NP and polymerase subunits. Immunostaining was performed after protease treatment of fixed and permeabilized cells. NP and PB2 (left), NP and PB1 (middle), NP and PA (right) were detected in the infected cells at 7 h post-infection (hpi). Scale bars, 10 µm. **c,** Nucleolar localization of vRNA. Negative-stranded vRNA of PB2 segment were detected in the infected cells at 5 hpi by fluorescence *in situ* hybridization. Nucleolin was immuno-stained. Insets: enlarged versions of the selected regions indicated by the white boxes. Scale bars, 10 µm. **d–i,** Immunogold-labelled ultrathin sections of mock-infected or PB2-FLAG virus-infected MDCK cells (MOI=5) at 5 hpi for protein detection: Yellow dashed circles mark normal nucleoli (**f, i**), relatively normal nucleoli (**d, g**), and abnormal nucleoli (**e, h**). Scale bars, 500 nm. FC, fibrillar centre; GC, granular component. **j,** Immunoprecipitation of vRNPs from the nucleoplasmic (Nu) and nucleolar (No) fractions of PB2-FLAG virus-infected MDCK cells (MOI=5) at 4 hpi. All images are representative of three independent experiments.

To further determine their exact localization in the nucleolus, MDCK cells were infected with a recombinant influenza A virus expressing C-terminally FLAG-tagged PB2 (PB2-FLAG virus) and subjected to immunoelectron microscopy. At 5 hpi, NPs were localized throughout the granular component (GC) regions, which are electron-dense areas involved in ribosome assembly, but not in fibrillar centre (FC) regions, where rRNA transcription occurs (Figures 1d and 1e). In contrast, although PB2 subunits were localized in the GC regions, they were mainly localized in periphery, but not central region, of the nucleolus (Figures 1g and 1h). These results suggest that the vRNP components are assembled into vRNP in the GC region of the nucleolar periphery.

To confirm that the nucleolus is the assembly site of vRNP components, we separated virus-infected cells into cytoplasmic, nucleoplasmic, and nucleolar fractions at 4 hpi; α-tubulin (cytoplasm marker), histone H3 (nucleoplasm marker), and nucleophosmin 1/B23 (NPM1, nucleolus marker) were detected in the expected fractions (Extended Data Figure 2). Then, PB2-FLAG was immunoprecipitated from the nucleoplasmic and nucleolar fractions, and the precipitates were examined. The co-precipitation of NP, PB1, and PA subunits with the PB2 subunit from the nucleoplasm fraction (Figure 1j) suggested that the vRNP components form vRNP complex. Likewise, these vRNP components were coprecipitated with PB2 from the nucleolar fraction. Collectively, our ultrastructural and biochemical data strongly suggest that the vRNP components are assembled to form vRNPs in the nucleolus.

### Importance of nucleolar NP localization for functional vRNP formation

Of the vRNP components, only NP possesses a nucleolar localization signal (NoLS) in addition to a nuclear localization signal^18,21,22^. To investigate the importance of NP nucleolar localization for the vRNP formation, we constructed mutant vRNPs using two NoLS-mutant NPs: NP^NoLSmut^ with alanine substitutions in the NoLS localizes only in the nucleoplasm (Extended Data Figures 3a and 3b) and a reverse mutant NoLS-NP^NoLSmut^, with an intact NoLS fused to the amino-terminus of NP^NoLSmut^ that causes its nucleolar localization (Extended Data Figures 3a and 3b)^18^. Strand-specific RT-qPCR after the plasmid-driven minigenome assay demonstrated that the vRNPs comprising NP^NoLSmut^ exhibited significant reduction in vRNA, cRNA, and mRNA production, while the vRNPs comprising NoLS-NP^NoLSmut^ showed relatively efficient production (Extended Data Figure 3c). These results indicate that the nucleolar localization of NP is critical for both transcription and replication of vRNA and are consistent with a previous report^18^, implying that the nucleolar localization of NP might be essential for functional vRNP formation.

To elucidate the impact of nucleolar NP localization on vRNP formation, we co-expressed PB2-FLAG, PB1, PA, and HA vRNA, together with wild-type NP (NP wt) or NP mutant, to reconstitute vRNPs in the cells. Then, the cells were subjected to immunoprecipitation using anti-FLAG antibody, and the precipitates were assessed by western blotting and RT-PCR (Figure 2a). NP wt, PB1, and PA were coprecipitated with PB2, and the full-length HA vRNA was also detected in the precipitate (Figure 2a). Additionally, the immunoprecipitated vRNPs produced HA mRNA by *in vitro* transcription (Figure 2b), indicating the assembly of these viral components into functional vRNPs. However, NP^NoLSmut^ was barely coprecipitated with PB2, although PB1 and PA were coprecipitated (Figure 2a). Furthermore, full-length HA vRNA was barely detected in the precipitate, and the immunoprecipitated vRNPs did not produce HA mRNA (Figure 2b), indicating that the NP^NoLSmut^ was not properly assembled into functional vRNPs, although heterotrimeric viral polymerase subunit was assembled. Intriguingly, NoLS-NP^NoLSmut^, PB1, and PA were adequately coprecipitated with PB2, from which full-length HA vRNA was detected (Figure 2a). Moreover, the immunoprecipitated vRNPs produced HA mRNA (Figure 2b), suggesting that some NoLS-NP^NoLSmut^ were assembled into functional vRNPs. Taken together, these results indicate that nucleolar localization of NP is indispensable for functional vRNP formation.

**Figure 2.**
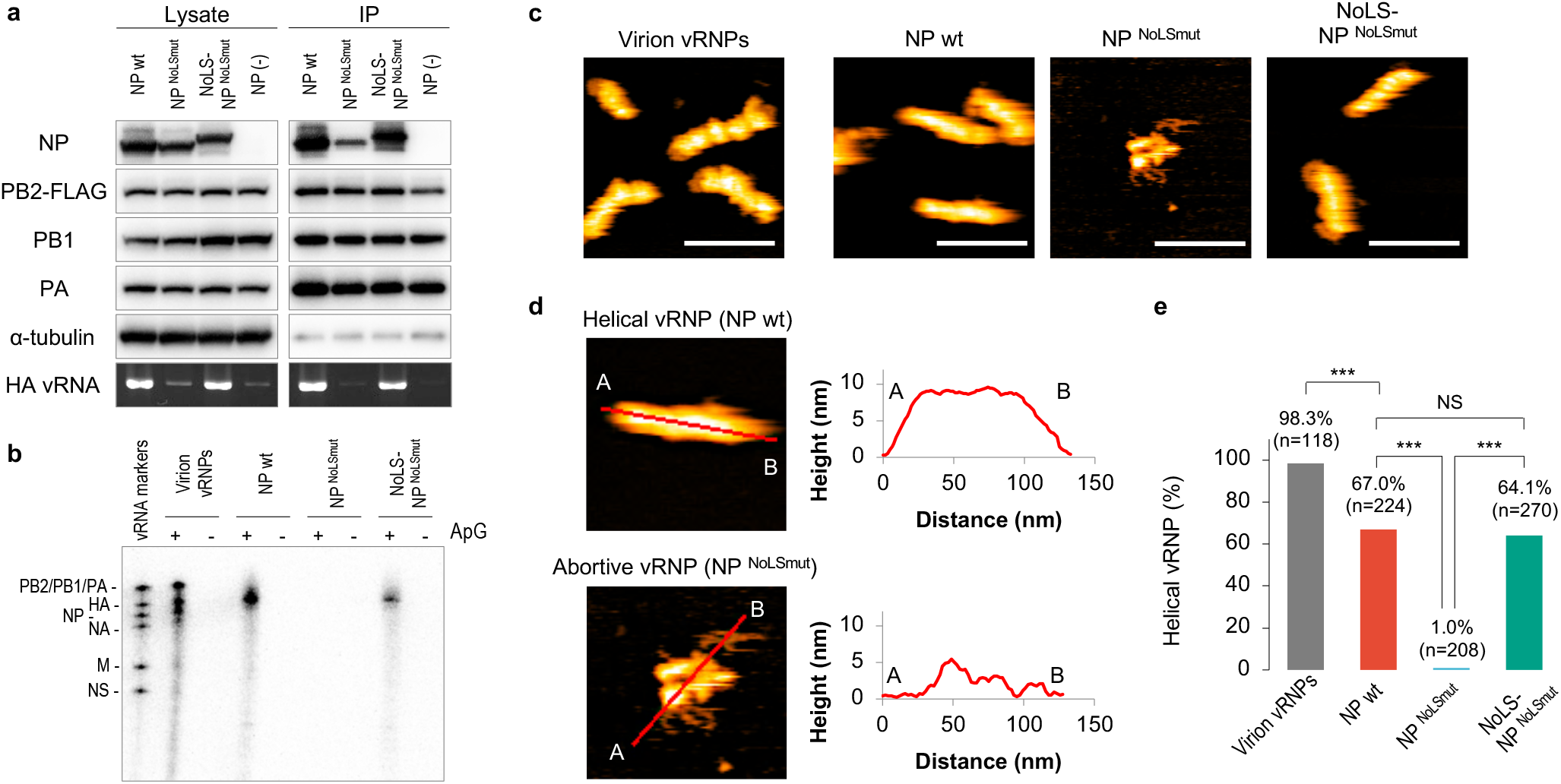
Nucleolar localization of NP is essential for helical vRNP formation. **a,** Reconstruction and immunoprecipitation of vRNPs. The vRNPs were reconstructed by transient expression of PB2-FLAG, PB1, PA, NP proteins and, HA vRNA in HEK293T cells, followed by immunoprecipitation using anti-FLAG antibody-conjugated agarose gels. The viral proteins and α-tubulin were immunoblotted. The full-length HA vRNA was detected by RT-PCR. Representative images from three independent experiments are shown. **b,** *In vitro* transcription of the reconstructed vRNPs. Nascent viral RNA was transcribed *in vitro* with ApG primer and detected by autoradiography. vRNPs derived from virion (virion vRNPs) were used as the control. vRNA markers were *in vitro* synthesized vRNAs by T7 RNA polymerase as the size markers. Representative images from three independent experiments are shown. **c,** HS-AFM observation of vRNPs. Representative images of the reconstructed vRNP and the virion vRNPs from two independent experiments are shown. Scale bars, 100 nm. **d,** Section analysis of the helical and abortive vRNPs. Left, enlarged HS-AFM images of Fig. 2c. Right, heights of the helical and the abortive vRNPs were measured at the red lines from A to B. **e,** Quantification of helical vRNP. The bars show the ratio of helical RNPs in all observed vRNPs in HS-AFM analysis. The ratio was compared using one-way ANOVA with Tukey test; \*\*\**P*<0.001, NS, not significant.

Ultrastructural analysis of the reconstituted vRNPs provided further evidence for the necessity of nucleolar NP localization for assembly into vRNPs. Using high speed atomic force microscopy (HS-AFM), which enables near-native topological ultrastructure visualization of biological specimens in solution without any fixation, hydration, and staining^23^, we investigated morphology of respective reconstituted vRNPs after immunoprecipitation and purification. Approximately 70% of the NP wt-constituted vRNPs showed double-helical structure with a uniform height of ∼9 nm (Figures 2c, 2d, 2e, and Supplementary movie 1). These vRNPs were morphologically indistinguishable from those purified from influenza virions (Figure 2c, Extended Data Figure 4b, and Supplementary movie 2). In contrast, NP^NoLSmut^ were barely assembled into double-helical structures and the resultant vRNPs showed pleomorphic morphology with a height of ≤5 nm, where string-like structures, probably naked RNAs based on their structure, were exposed (Figure 2c, 2d, 2e, and Supplementary movie 3). Importantly, NoLS-NP^NoLSmut^ was also assembled into double-helical vRNPs (∼65% of the vRNPs) (Figure 2c, 2d, 2e, and Supplementary movie 4). Immunoelectron microscopy confirmed that both double-helical vRNPs and the pleomorphic aggregates comprised NP and viral polymerase (Extended Data Figure 4a), indicating that the pleomorphic aggregates are abortive vRNPs. Taken together, these data demonstrate that the nucleolar NP localization is critical for functional double-helical vRNP formation.

### Impact of nucleolar disruption on functional vRNP formation

Considering the necessity of nucleolar NP localization for proper vRNP formation, nucleolar structure disruption would heavily impact the vRNP component assembly. To test this hypothesis, we used a selective RNA polymerase I (Pol I) inhibitor, CX5461^24^; inhibition of Pol I activity that transcribes 47S ribosomal RNA (pre-rRNA) causes translocation of some nucleolar proteins to the nucleoplasm, resulting in nucleolar disruption^25,26^. Actinomycin D, which inhibits both Pol I and Pol II activities, was used as control. RT-qPCR revealed that CX5461 treatment (2–10 μM) inhibited only pre-rRNA transcription (Figure 3a and Extended Data Figure 5a), whereas actinomycin D treatment (10 μg/mL) suppressed the transcription of both pre-rRNA (Pol I) and pre-mRNA (Pol II) (Figure 3a), indicating that CX5461 treatment specifically inhibits Pol I activity. In addition to an rRNA staining dye, immunostaining using an antibody against nucleolin, a nucleolar marker localized in GC region, showed that nucleolin in CX5461-treated cells was translocated from the nucleolus to the nucleoplasm in a dose-dependent manner and that the morphology of the pleomorphic nucleoli was altered into small spherules (Figure 3b), demonstrating that CX5461 caused nucleolar disruption through Pol I activity inhibition.

**Figure 3.**
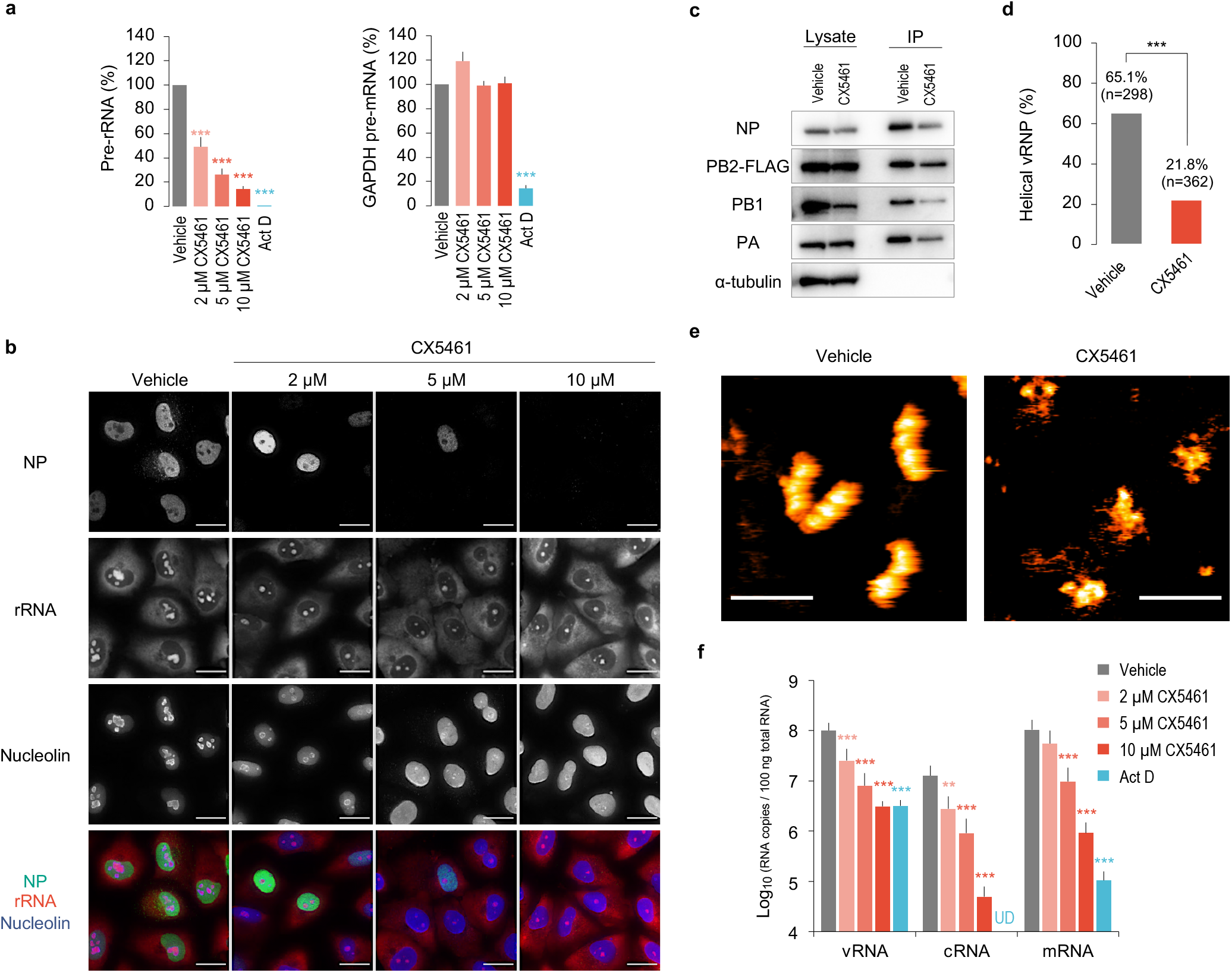
Nucleolar disruption induced by an RNA polymerase I inhibitor impairs viral replication, transcription, and helical vRNP formation. **a, b, f,** A549 cells were pretreated with CX5461, 10 µg/mL actinomycin D (Act D), or 1% DMSO (Vehicle) for 2 h, followed by wild-type virus infection (MOI=5) for 5 h. **a,** Selectivity of the RNA polymerase inhibitors on Pol I and II activities. Total RNA was extracted and analysed by RT-qPCR. The expression levels were compared with that of vehicle-treated cells using one-way ANOVA with Dunnett’s test; \*\*\**P*<0.001. Data are presented as mean ± S.D. of three independent experiments. **b,** CX5461-induced nucleolar disruption and its effect on NP expression. Scale bars, 20 µm. Representative images from three independent experiments. **c,** Immunoprecipitation of vRNPs from the PB2-FLAG virus-infected A549 cells (MOI=5), followed by 10 µM CX5461 or vehicle treatment at 2 hpi. The cells were lysed at 4.5 hpi and immunoprecipitated. Representative images from three independent experiments. **d,** Quantification of helical vRNP. The bars show the ratio of helical to total vRNPs in HS-AFM analysis. **e,** Representative images of the vRNPs in HS-AFM analysis. Scale bars, 100 nm. The ratio was compared using Welch t-test; \*\*\**P*<0.001. **f,** Effects of the nucleolar disruption on viral replication and transcription. HA vRNA, cRNA, and mRNA copy numbers were measured by strand-specific RT-qPCR and compared with that of vehicle-treated cells using one-way ANOVA with Dunnett’s test; \*\**P*<0.01, \*\*\**P*<0.001, UD, undetected. Data are presented as mean±S.D. of three independent experiments with two RT-qPCR assays.

To determine whether nucleolar disruption affects vRNP formation, PB2-FLAG virus-infected A549 cells were treated with 10 μM CX5461 at 2 hpi, and vRNPs were immunoprecipitated using anti-FLAG antibody at 4.5 hpi (Figure 3c and Extended Data Figure 6a). CX5461 treatment modestly decreased the amount of immunoprecipitated NP as well as PB1 and PA subunits in these cells, although viral protein expression levels were comparable, or marginally lower, compared to those in control cells (Figure 3c), suggesting that nucleolar disruption impacted the vRNP component assembly. Importantly, ultrastructural analysis of the immunoprecipitated and purified vRNPs using HS-AFM revealed a significant reduction in efficiency of double-helical vRNP formation in CX5461-treated cells (Figure 3d). Most of the vRNPs immunoprecipitated from CX5461-treated cells were pleomorphic aggregates (Figure 3e and Extended Data Figures 6b and 6c) that were similar to the abortive vRNPs composed of NP^NoLSmut^ (Figure 2c), while most vRNPs immunoprecipitated from control cells had double-helical structures (Figure 3d). Consistent with the ultrastructural analysis, HA vRNA, cRNA, and mRNA production (Figure 3f) and viral growth (Extended Data Figures 5b and 5c) were decreased in CX5461-treated cells in a dose-dependent manner, without any cell toxicity (Extended Data Figure 5d). Thus, these results demonstrate that the nucleolus is required for proper assembly of the vRNP components into functional double-helical vRNPs.

## Discussion

We showed that the nucleolus is the site for formation of functional vRNPs with double-helical structure. At an early infection stage, all vRNP components were localized in the nucleolus. Inhibition of nucleolar NP localization and nucleolar structure disruption affected vRNP component assembly, resulting in defective vRNP formation. These results demonstrated that the vRNP components are assembled into double-helical vRNPs in the nucleolus and that the nucleolus plays an important role in influenza virus genome replication.

Using two NP mutants, we showed that nucleolar NP localization via NoLS is required for functional vRNP formation (Figure 2), indicating that NP plays a pivotal role in vRNP formation in the nucleolus. Then, how is the heterotrimeric viral polymerase translocated to the nucleolus in absence of NoLS? One possibility is that it is transported in association with NS1, which is a non-structural viral protein and is translocated to the nucleolus via its NoLS^27,28^. However, because NS1 unlikely interacts with viral polymerase subunits^29^ and is not essential for vRNP formation as well as vRNA transcription and replication (Figure 2), NS1 would not be involved in viral polymerase transport to the nucleolus. Because NP interacts with PB2 and PB1 subunits^30^, it would be responsible for viral polymerase import into the nucleolus. Indeed, although mutations in NP residues that are required for interaction with viral polymerase do not affect its properties, namely, nuclear localization, RNA binding, and oligomerization, they significantly impact vRNA transcription and replication^31^, suggesting that the viral polymerase might not be transported into the nucleolus by the mutant NP, resulting in incomplete vRNP formation. Thus, interaction between viral polymerase and NP is likely involved in its nucleolar import and subsequent vRNP formation.

Nucleolar disruption by a specific Pol I inhibitor disrupted vRNP component assembly into functional double-helical vRNPs (Figure 3). Since Pol I-mediated pre-rRNA transcription is required for nucleolar structure maintenance^25,26^, it is possible that certain nucleolar proteins, which are required for vRNA replication-coupled vRNP formation, were translocated outside the nucleolus by the Pol I inhibitor treatment. Several host nucleolar proteins, including nucleolin and NPM1, reportedly interact with NP, and some nucleolar proteins, such as nucleolin, NPM1, LYAR, and FMRP, facilitate vRNA replication and vRNP assembly^14,16, 32–34^. Thus, Pol I activity inhibition would change their localizations and disrupt their proper interactions with NP in the nucleolus, resulting in abortive vRNP formation.

Several studies imply the involvement of the nucleolus in vRNA replication. Khatchikian *et al.* reported that host 28S rRNA-derived 54 nucleotides are inserted into the HA vRNA during viral replication via genetic recombination^35^. This recombination is probably caused by polymerase jumping mechanism^35,36^, wherein the viral polymerase transitions between HA vRNA and an adjacent host 28S rRNA during vRNA replication, suggesting that the replication occurs at the site of rRNA transcription or at its adjacent site, i.e., the FC region in the nucleolus. Subsequently, vRNP assembly occurs in the GC region (Figure 1d-1), where ribosome assembly occurs. Accordingly, an *in situ* hybridization study on salmon anaemia virus-infected cells (also belonging to *Orthomyxoviridae* family) showed nucleolar localization of anti-genomic as well as genomic RNA^37^. Although identity of the anti-genomic RNA in the nucleolus remains uncertain, considering that viral mRNA is transcribed in the vicinity of Pol II in the nucleoplasm, the anti-genomic RNA likely represents cRNA replicated from vRNA template. Moreover, we detected not only vRNA (Figure 1c) but also anti-genomic RNA (Extended Data Figure 7) in the nucleolus of virus-infected cells, supporting that the nucleolus is the site of vRNA replication and vRNP formation.

In conclusion, we demonstrated that the formation of functional double-helical vRNP occurs in the nucleolus. Our results highlight the importance of the nucleolus during influenza virus life cycle. Further studies on intra-nucleolar host factors responsible for vRNP formation are necessary to understand the detailed mechanisms of vRNP formation, which would contribute to the development of novel antivirals against influenza viruses.

## Methods

### Cell culture

Madin–Darby canine kidney (MDCK) cells were grown in minimal essential medium (MEM) (Thermo Fisher Scientific, MA USA) containing 5% newborn calf serum (16010-159, Thermo Fisher Scientific). Human embryonic kidney 293T (HEK293T) cells (CRL-3216) and human lung carcinoma (A549) cells (CCL-185) were purchased from ATCC (Manassas, VA USA) and maintained in Dulbecco’s modified Eagle medium (Merck, Germany) supplemented with 10% foetal bovine serum (FB-1365, Biosera, France). Cultures were maintained at 37 °C in a 5% CO_2_ atmosphere. Viruses were grown in MEM containing 0.3% bovine serum albumin (BSA/MEM).

### Plasmid construction

pCAGGS/NP^NoLSmut^ and pCAGGS/NoLS-NP^NoLSmut^ were constructed using inverse PCR^38^ with sequences similar to those previously reported (pCAGGS/NP-NLS2mut and pCAGGS/NLS2-NP-NLS2mut, respectively)^18^. To generate pCAGGS/PB2-FLAG, the PB2 ORF and FLAG (DYKDDDDK) were linked with a linker (AAA). pPol I/PB2-FLAG was constructed by inserting the PB2-FLAG ORF with stop codon into a truncated pPol I/PB2 plasmid with 3′ non-coding region and additional 143 nucleotides of 5′ terminal coding and non-coding regions^39^.

### Inhibitors and antibodies

Inhibitors used were: CX5461 (CS-0568, ChemScene, NJ USA), actinomycin D (A1410, Merck), and cycloheximide (037-20991, Fujifilm, Japan). The primary antibodies used for immunofluorescence, western blotting, and immuno-electron microscopy were: anti-NP mouse monoclonal^40^, anti-NP rabbit polyclonal (GTX125989, GeneTex, CA USA), anti-PB2 goat polyclonal (sc-17603, Santa Cruz Biotechnology, TX USA), anti-PB1 goat polyclonal (sc-17601, Santa Cruz Biotechnology), anti-PA rabbit polyclonal (GTX125932, GeneTex), anti-nucleolin rabbit polyclonal (ab22758, Abcam, UK), anti-nucleophosmin mouse monoclonal (ab10530, Abcam), anti-α-tubulin rabbit polyclonal (PM054, Medical & Biological Laboratories, Japan), anti-histone H3 rabbit polyclonal (GTX122148, GeneTex), anti-digoxigenin sheep polyclonal (11 333 089 001, Roche, Switzerland), and anti-FLAG mouse monoclonal (M185-A48, Medical & Biological Laboratories). The secondary antibodies used were: Alexa fluor 488-conjugated anti-mouse (A11001, Thermo Fisher Scientific), anti-rabbit (A11008, Thermo Fisher Scientific), anti-goat (A11055, Thermo Fisher Scientific), anti-sheep (A11015, Thermo Fisher Scientific), Alexa555-conjugated anti-mouse (A21422, Thermo Fisher Scientific), anti-rabbit (A21428, Thermo Fisher Scientific), Alexa405-conjugated anti-rabbit (ab175652, Abcam), HRP-conjugated anti-mouse (NA931, GE Healthcare, IL USA), anti-rabbit (NA934, GE Healthcare), anti-goat (ab6741, Abcam), 6 nm gold-conjugated anti-mouse (715-195-150, Jackson ImmunoResearch, PA USA), and anti-rabbit (711-195-152, Jackson ImmunoResearch).

### Generation of recombinant viruses by reverse genetics

Reverse genetics was performed using pPol I plasmids containing cDNA sequences of the A/WSN/33 (WSN; H1N1) viral genes between the human Pol I promoter and mouse Pol I terminator as described previously^41^. Briefly, eight pPol I plasmids and pCAGGS protein-expression plasmids for PB2, PB1, PA, and NP were mixed with TransIT-293 (Mirus Bio, WI USA) and added to HEK293T cells. Forty-eight hours post-transfection, the cells were treated with 1 μg/mL TPCK-Trypsin (Worthington, OH USA) for 30 min, centrifuged at 1,750 × g for 15 min at 4 °C, and the supernatant was collected and stored at −80 °C. PB2-FLAG virus was generated by replacing pPol I/PB2 wt with pPol I/PB2-FLAG plasmid. For subsequent viral amplification, MDCK cells were infected at MOI of 10^-5^ and incubated for two days in BSA/MEM containing 1 μg/mL TPCK-Trypsin. Viral titres were determined by plaque assay using MDCK cells.

### Immunofluorescence

Cells were plated in 8-well chamber slides (Matsunami, Japan) coated with rat collagen I (Corning, NY USA). Infected or transfected cells were fixed in 4% paraformaldehyde (PFA) in phosphate buffer (PB) (Nacalai Tesque, Japan) for 10 min and then permeabilized with 0.5% Triton-X in PBS for 5 min. The cells were blocked with Blocking One (Nacalai Tesque) for 30 min followed by incubation with primary antibodies overnight at 4 °C and secondary antibodies for 1 h at room temperature. For nuclei and rRNA staining, cells were treated with Hoechst 33342 (Thermo Fisher Scientific) and Nucleolus Bright Red (Dojindo, Japan), respectively, for 10 min. Section images were recorded using DeltaVision Elite (GE healthcare) with a 60× oil objective, deconvolved and projected using ‘Quick Projection’ tool by softWoRx (GE Healthcare).

### Protease treatment

As the optimal condition for protease treatment depends on the protease type, lot, and cell strain^20^, we recommend verifying the protease concentration and incubation time. After permeabilization, the cells were washed twice in cold-PBS on ice and placed in cold 5 µg/mL TPCK-Trypsin in PBS. The slides were incubated on a plate incubator (MyBL-P2, AS ONE, Japan) at 37 °C for 5 min and incubated with cold 4% PFA in PB (final concentration 2%) on ice for 30 min to terminate reaction. Thereafter, the cells were washed in PBS and blocked as described above.

### Fluorescence *in situ* hybridisation (FISH)

FISH was performed as described previously^42^. Briefly, probes were transcribed *in vitro* using digoxigenin (DIG)-11-UTP (Roche) and RiboMAX Large Scale RNA Production System-T7 (Promega, WI USA). The template of positive- and negative-sense PB2 genome segment (∼300 bp) was PCR amplified using pPol I/PB2. The primers used are listed in Supplementary Table 1.

The infected cells were fixed with 4% PFA in PB for 10 min and permeabilized with 0.5% Triton X-100 for 5 min at room temperature. Subsequently, cells were sequentially washed with 2× and 0.01× SSC (Nacalai Tesque), incubated in 95% formamide in 0.1× SSC for 15 min at 65 °C, and immediately chilled on ice. Cells were then blocked with prehybridization buffer (50% formamide [Fujifilm], 2× SSC, 5× Denhardt’s solution [Fujifilm], 20 μg/mL salmon sperm DNA [BioDynamics Laboratory, Japan]) for 1 h at room temperature and then incubated with 200 ng/mL of DIG-labelled RNA probe diluted in prehybridization buffer overnight at 60 °C on a shaker. After hybridization, cells were thoroughly washed with wash solution 1 (50% formamide, 2× SSC, 0.01% Tween-20) and wash solution 2 (0.1× SSC, 0.01% Tween-20) (three washes with each buffer for 20 min/wash at 60 °C). Finally, cells were incubated with *in situ* hybridization blocking solution (Vector Laboratories, CA USA) for 30 min at room temperature, and probes were detected by immunofluorescence using anti-DIG sheep and Alexa fluor488-conjugated anti-sheep antibodies.

### Western blotting

Western blotting was performed as previously described^17^. Briefly, cells or samples described below were dissolved with 2× Tris-Glycine SDS Sample Buffer (Thermo Fisher Scientific), boiled for 5 min in absence of a reducing agent, and subjected to SDS-PAGE. Proteins were electroblotted onto Immobilon-P transfer membranes (Merck). The membranes were blocked with Blocking One for 30 min at room temperature and then incubated with primary antibodies overnight at 4 °C. After incubation with HRP-conjugated secondary antibodies for 1 h at room temperature, the blots were developed using Chemi-Lumi One Super (Nacalai Tesque).

### Cell viability

Cell viability was assessed with CellTiter-Glo Luminescent Cell Viability Assay (Promega) according to the manufacturer’s instructions. Briefly, CellTiter-Glo reagent (equal in volume to the culture medium) was added to A549 cells. Plates were shaken on a plate shaker for 2 min to induce cell lysis, incubated at room temperature for 10 min, and subjected to luminescence measurement.

### vRNP reconstruction and immunoprecipitation

HEK293T cells were plated in two 10 cm^2^ dishes and transfected using PEI MAX (Polysciences, PA USA) with vRNP expression plasmids (3 µg/mL each of pCAGGS/PB2-FLAG, pCAGGS/PB1, pCAGGS/PA, and pCAGGS/NP; 300 ng/µL pPol I/HA). Two days post-transfection, cells were suspended in cold-PBS and pelleted by centrifugation at 780 × g for 10 min at 4 °C. The pellets were resuspended in 500 µL lysis buffer (50 mM Tris-HCl pH 8.0, 150 mM NaCl, 5 mM MgCl_2_, 10% Glycerol, 0.05% NP-40, 2 mM DTT, 10 mM Ribonucleoside-Vanadyl complex [New England Biolabs, MA USA], 1× Protease inhibitor complete EDTA-free [Roche]), rotated for 15 min at 4 °C, and centrifuged at 20,000 × g for 15 min at 4 °C. The pellets were resuspended in the buffer and incubated with additional 80 µL anti-FLAG M2 affinity gel (Merck) on a rotator overnight at 4 °C. The gels were washed once with lysis buffer, thrice with wash buffer (50 mM Tris-HCl pH 8.0, 200 mM NaCl, 50 mM Na_2_HPO_4_, 2 mM DTT), and eluted in 150 µL wash buffer with 500 ng/µL FLAG peptide (Merck) by rotation with a rotator for 30 min at 4 °C. Cell lysates and eluates were electrophoresed with SDS-polyacrylamide gel and immunoblotted.

### vRNP purification

vRNP purification was performed as described previously^43^. To prepare virion-derived vRNPs, MDCK cells were infected with the virus and incubated at 37 °C for two days. Virions in the supernatants were purified by ultracentrifugation through a 30% (w/w) sucrose cushion. The pellets were resuspended in PBS. The purified virions were lysed in a solution containing 50 mM Tris-HCl pH 8.0, 100 mM KCl, 5 mM MgCl_2_, 1 mM DTT, 2% lysolecithin, 2% Triton X-100, 5% glycerol, and 1 U/μL RNase inhibitor (Promega) for 1 h at 30 °C.

The lysed or immunoprecipitated vRNPs were ultracentrifuged through a glycerol gradient (30%–70%) containing 50 mM Tris-HCl pH 8.0 and 150 mM NaCl at 245,000 × g for 3 h at 4 °C. Each fraction was electrophoresed with SDS-polyacrylamide gel and immunoblotted with an anti-NP antibody (Supplementary Figure 1 and Extended Data Figure 6a). NP-enriched fractions 7 and 8 were used for vRNP observations.

### *In vitro* transcription of vRNPs

The purified vRNP (0.01 mg/mL) was incubated in a buffer (50 mM Tris-HCl buffer pH 7.9, 5 mM MgCl_2_, 40 mM KCl, 1 mM DTT, 10 μg/mL actinomycin D, 1 mM each of ATP, CTP, and GTP, 0.25 μCi/μL [α-^32^P] UTP and 0.05 mM UTP, 1 U/μL RNasin Plus RNase inhibitor, 1 mM ApG [IBA, Germany]) at 30 °C for 15 min. RNA was purified using RNeasy Mini kit, mixed with equal volume of 2× RNA Loading Dye (New England Biolabs), heated at 90 °C for 2 min, and immediately chilled on ice. The sample was electrophoresed on 4% polyacrylamide gel containing 7 M urea in 0.5× TBE buffer (Nacalai Tesque) at 120 V for 5 h. The gel was dried at 80 °C for 2 h, exposed to an imaging plate (BAS-MS 2025, Fujifilm) for 12–24 h, and scanned with a Typhoon 3000 Phosphorimager (GE Healthcare). For preparation of vRNA markers, all eight vRNA segments of the influenza virus were transcribed using 0.25 μCi/μL [α-^32^P] UTP and RiboMAX Large Scale RNA Production System-T7 as described above. The transcribed RNAs were purified and mixed before electrophoresis.

### High-speed atomic force microscopy (HS-AFM)

HS-AFM analysis of vRNP was performed as described by Nakano *et al*.^44^. The samples were prepared in a microcentrifuge tube, dropped onto freshly cleaved mica without any surface modification, and incubated for 1–5 min at room temperature (∼24°C). The samples on the mica surface were then washed with imaging buffer (50 mM Tris-HCl pH 7.9, 5 mM MgCl_2_, 40 mM KCl, 1 mM DTT), and observed in the imaging buffer at room temperature (∼24°C) using High-Speed Atomic Force Microscope SS-NEX (RIBM, Japan). Images were taken at a 2 images/sec rate using cantilevers (BL-AC10DS, Olympus, Japan) with a 0.1 N/m spring constant and a resonance frequency in water of 0.6 MHz. To increase the resolution, the electron-beam deposited tips were fabricated using phenol or ferrocene powder^45^. All HS-AFM images were viewed and analysed using Kodec software (version 4.4.7.39)^46^. A low-pass filter and a flattening filter were applied to individual images to remove spike noise and flatten the xy-plane, respectively. Rod-like and helical structures with a uniform height of 9.0 ± 1.5 nm were defined as helical vRNPs. Pleomorphic nucleic acid-protein aggregates, except for nucleic acids (<2.5 nm height string-like structures) or proteins (<25 nm long globular structures), were defined as abortive vRNPs.

### Immuno-electron microscopy

Purified vRNPs were adsorbed onto carbon-coated nickel grids and fixed with 2% PFA for 5 min. The grids were washed, treated with Blocking One, and then incubated with an anti-NP or anti-FLAG antibody overnight at 4 °C or for 1 h at room temperature, respectively. After washing, the grids were incubated with 6-nm gold conjugated anti-mouse or anti-rabbit antibodies for 1 h at room temperature. After washing, the samples were fixed with 2% PFA for 10 min and negatively stained with 2% uranyl acetate solution. The images were acquired with an HT7700 (Hitachi High-Tech Corporation, Japan).

For thin-section preparations, infected and mock-infected MDCK cells were fixed with 1.5% PFA and 0.025% glutaraldehyde in 0.1 M PB for 1 h. The fixed cells were dehydrated in a series of ethanol gradient and then embedded in LR-White resin. Ultrathin sections (80 nm) were cut with Leica EM UC7 (Leica, Germany) and collected on a nickel grid. Immuno-labelling was performed as described above without post-staining.

### RT-PCR

Total RNAs were extracted using an RNeasy Mini Kit with on-column DNase digestion (Qiagen). Ten nanograms of the extracted RNA samples were reverse-transcribed using a Uni-12 primer (5′-AGCRAAAGCAGG-3′) and Superscript III reverse transcriptase (Thermo Fisher Scientific). Ten-fold diluted cDNAs were PCR amplified using KOD FX (Toyobo, Japan) and 0.25 μM HA segment-specific primers according to manufacturer’s protocol. Cycling conditions were: initial denaturation at 2 min at 94 °C, followed by 25 cycles of 98 °C for 10 s, 55 °C for 30 s, and 68 °C for 2 min. The PCR products were electrophoresed on 1.0% agarose gels containing 0.01% (w/v) ethidium bromide in 0.5× TBE. The primers used are listed in Supplementary Table 2.

### RT-qPCR

Two-hundred nanograms of total RNAs were reverse-transcribed using Random primer 6 (New England Biolabs) and Superscript III reverse transcriptase. For qPCR, reactions contained 1 μL 10-fold diluted RT product, 7.5 μL THUNDERBIRD SYBR qPCR Mix, and 0.25 μM primers, at a final volume of 15 μL. Cycling conditions were: initial denaturation for 2 min at 94 °C, followed by 40 cycles of 98 °C for 10 s, 55 °C for 15 s, and 72 °C for 30 s. The relative expression levels of target genes were normalized to that of GAPDH. The primers used are listed in Supplementary Table 2. A primer set for pre-rRNA, described previously^47^, was used.

### Strand-specific RT-qPCR

Strand-specific RT-qPCR was performed as described previously^48,49^. Briefly, total RNA was extracted from cells using an RNeasy Mini Kit. cDNAs complementary to the three types of HA genome were synthesized with tagged primers at the 5′ end. A 2.5 μL mixture containing the 200 ng total RNA sample and 20 pmol tagged primers was heated for 10 min at 65 °C, chilled immediately on ice for 5 min, and then reheated to 60 °C. After 5 min, 7.5 μL of preheated reaction mixture [2 μL 5× First Strand buffer, 0.5 μL 0.1 M dithiothreitol, 0.5 μL dNTP mix (10 mM each), 0.5 μL Superscript III reverse transcriptase (200 U/μL), 0.25 μL RNasin Plus RNase inhibitor (40 U/μL, Promega), and 3.75 μL saturated trehalose] was added and incubated at 60°C for 1 h. For qPCR, each 15-microliter reaction contained 1 μL 50-fold diluted RT product, 7.5 μL THUNDERBIRD SYBR qPCR Mix, and 0.25 μM primers. Cycling conditions were: initial denaturation for 2 min at 95 °C, followed by 40 cycles of 95 °C for 10 s and 60 °C for 45 s. Ten-fold serial dilutions (10^9^,10^8^,10^7^,10^6^,10^5^,10^4^ copies/μL) of synthetic vRNA standards were used to generate a standard curve. The primers used are listed in Supplementary Table 3.

### Subcellular fractionation

We performed subcellular fractionation as described previously^50^ and optimized the buffers, incubation time, and centrifugal force for MDCK cells. Briefly, pelleted MDCK cells (two 15 cm^2^ dishes) were resuspended in ice-cold mild detergent buffer (20 mM Tris pH 7.5, 10 mM KCl, 3 mM MgCl_2_, 0.1% NP40, 10% glycerol) and centrifuged at 100 × g for 5 min at 4 °C. The supernatants were further centrifuged at 1,400 × g for 10 min at 4 °C and collected as the cytoplasmic fraction. Pellets were then resuspended in 3 mL of 0.25 M sucrose/10 mM MgCl_2_, layered over a 3 mL cushion of 0.35 M sucrose/3 mM MgCl_2_ and centrifuged at 1,400 × g for 5 min at 4 °C. The resulting cleaner nuclear pellet was resuspended in 0.35 M sucrose/3 mM MgCl_2_ and sonicated six times for 10 s on ice (10-second rest between pulses) to disrupt nuclei and release nucleoli. The sonicate was layered over a 3 mL cushion of 0.88 M sucrose/3 mM MgCl_2_ and centrifuged at 2,800 × g for 10 min at 4 °C to pellet nucleoli and the supernatant was collected as the nucleoplasmic fraction. All solutions used in the fractionation were supplemented with Protease inhibitor complete EDTA-free to minimize protein degradation.

Nucleoli were washed by resuspending in 0.5 mL of 0.35 M sucrose/3 mM MgCl_2_ followed by centrifuging at 2,800 × g for 5 min at 4 °C. The nucleolar pellet was resuspended in 300 μL middle salt RIPA buffer (50 mM Tris pH 7.5, 300 mM NaCl, 1% NP-40, 0.5% deoxycholate, Protease inhibitor complete EDTA-free) containing 16 μL of 1 unit/μL RQ1 RNase-free DNase and rotated for 30 min at 4 °C. The lysate was centrifuged at 20,000 × g for 10 min at 4 °C, the supernatant collected as the nucleolar extract, and the NaCl concentration adjusted to 150 mM by adding 300 μL of ‘no salt’ RIPA buffer (50 mM Tris pH 7.5, 1% NP-40, 0.5% deoxycholate, Protease inhibitor complete EDTA-free).

The cytoplasmic and nucleoplasmic fractions were mixed in 1× RIPA buffer (50 mM Tris pH 7.5, 150 mM NaCl, 1% NP-40, 0.5% deoxycholate, Protease inhibitor complete EDTA-free) and centrifuged at 2,800 × g for 10 min at 4 °C. Total protein concentrations were measured using Pierce BCA Protein Assay Kit (Thermo Fisher Scientific) and adjusted to approximately 0.5 µg/mL. The samples (∼0.5 µg) were subjected to western blotting.

### Statistics

We compared group means by Welch t-test or one-way analysis of variance (ANOVA) with Dunnett’s test, Tukey test, or two-way ANOVA, comparing each group with the indicated control using R packages^51^. We considered a P value ≤ 0.05 to be statistically significant.

### Data availability

All data are available from the corresponding author upon request. Source data for gels and blots are provided as Supplementary Information.

## Acknowledgements

We thank Y. Kawaoka for providing plasmids for the generation of influenza A virus WSN strain, F. Momose for providing mAb61A5, N. Kodera for preparing cantilevers for HS-AFM analysis, and K. Shindo for her help with vRNP reconstruction. This work was supported by JSPS KAKENHI Grant 19J14928 (to S.M.), JSPS Grant-in-Aid for Scientific Research (B) (17H04082, 20H03494), JSPS Grant-in-Aid for Challenging Research (Exploratory) (19K22529), the JSPS Core-to-Core Program A, the Advanced Research Networks, MEXT Grant-in-Aid for Scientific Research on Innovative Area (19H04831), an AMED Research Program on Emerging and Re-emerging Infectious Disease grants (19fk0108113, 20fk0108270h0001), the JST Core Research for Evolutional Science and Technology, the Grant for Joint Research Project of the Institute of Medical Science, University of Tokyo, the Joint Usage/Research Center program of Institute for Frontier Life and Medical Sciences Kyoto University, the Daiichi Sankyo Foundation of Life Science, the Uehara Memorial Foundation, and the Takeda Science Foundation (to T.N.).

## Author contributions

S.M., M.N., and T.N. designed the study; S.M., M.N., T.M., A.H., R.T., Y.F., and N.H. performed experiments; S.M., M.N., Y.M., and T.N. wrote the manuscript with input from all co-authors.

## Competing interest declaration

The authors declare no competing interests.

## Extended data figure legends

**Extended Data Figure 1.**
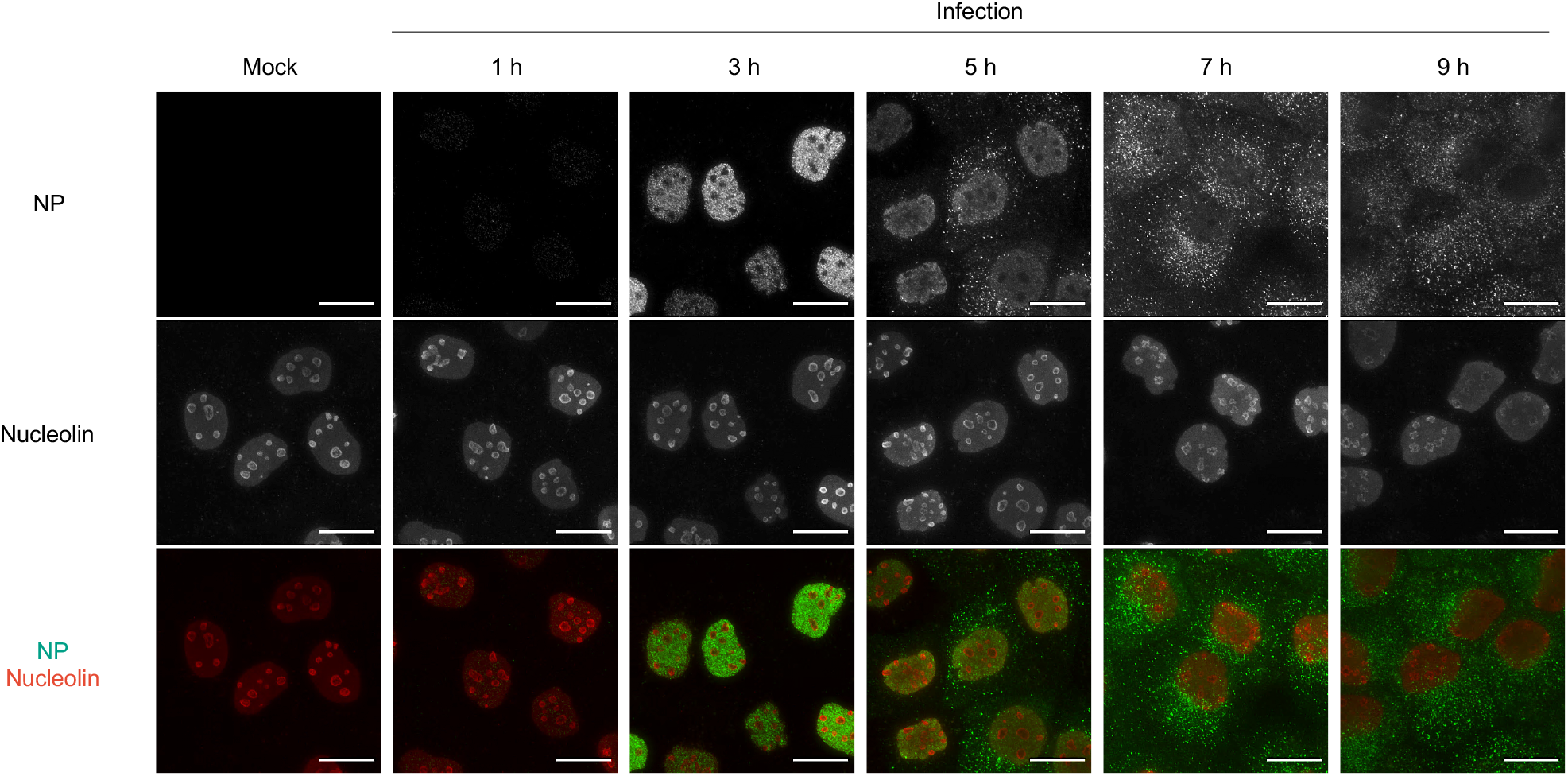
Immunofluorescence assay without protease treatment in influenza virus-infected cells. **a,** Subcellular translocation of NP in the infected cells. This experiment was performed in parallel that depicted in Figure 1a, and its fluorescence signals were unified. NP and nucleolin were immuno-stained without protease treatment. Scale bars, 20 µm. Representative images from three independent experiments.

**Extended Data Figure 2.**
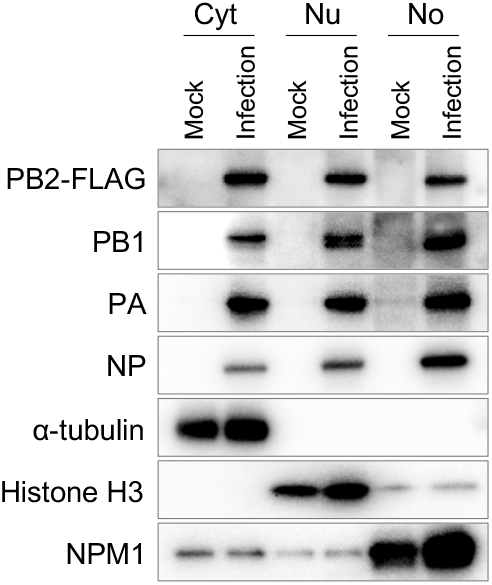
Subcellular fractionation of the infected cells. The mock-infected or PB2-FLAG virus-infected MDCK cells at an MOI of 5 were fractionated into cytoplasmic (Cyt), nucleoplasmic (Nu), and nucleolus (No) fractions at 4 hpi. Approximately 5 µg total protein were analysed by western blotting of viral proteins and cell fraction-specific markers α-tubulin (Cyt), histone H3 (Nu), and NPM1 (No). Representative images from three independent experiments.

**Extended Data Figure 3.**
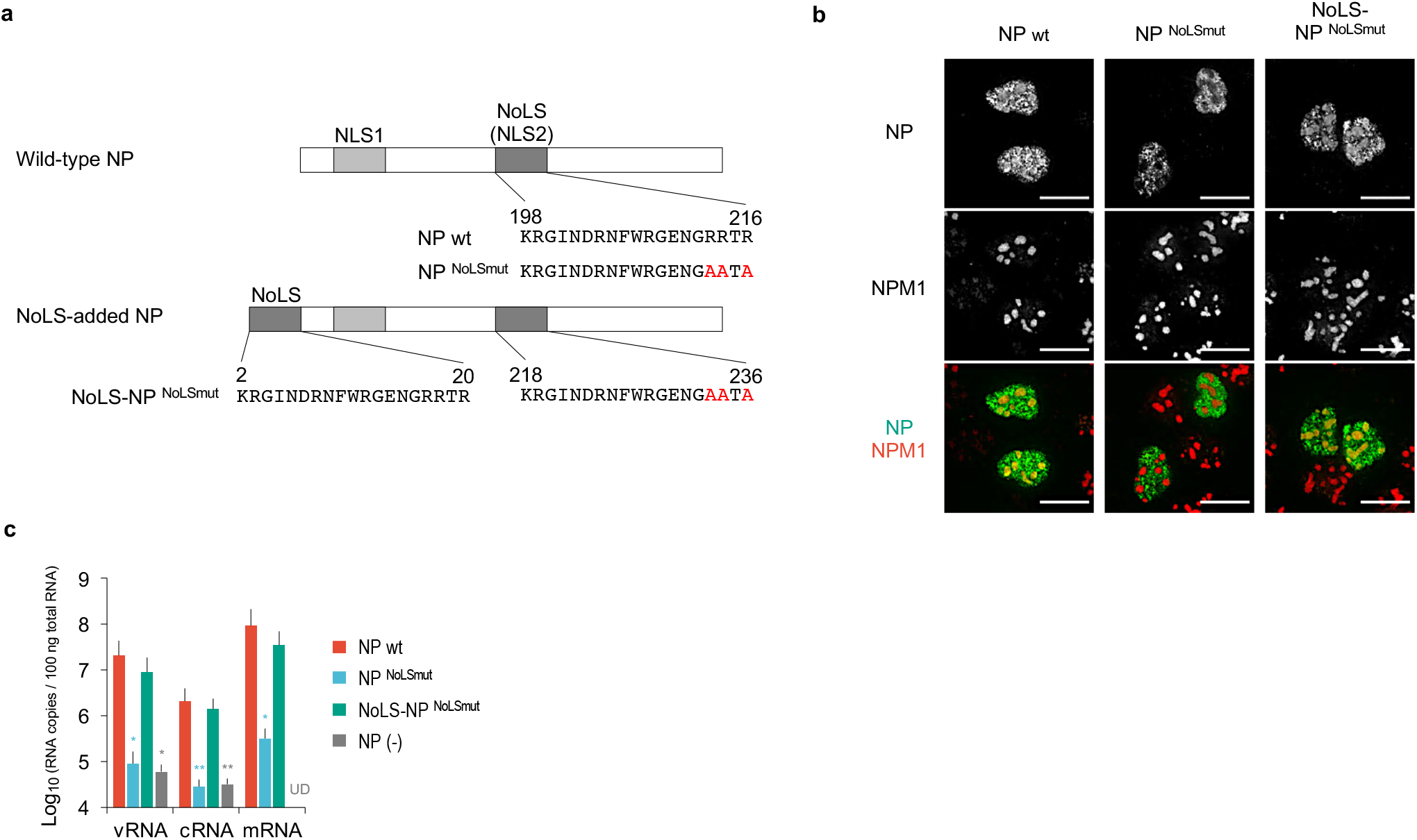
Nucleolar localization of NP is critical for transcription and replication. **a,** Schematic diagram of NP wt, NP^NoLSmut^, and NoLS-NP^NoLSmut^. NLS1 (residues 3 to 13) and NoLS (NLS2, residues 198 to 216) are represented in light and dark grey colour, respectively. The NoLS motif was added to the amino terminus of NP^NoLSmut^. The alanine replacements are in red. **b,** Nucleolar localization of the overexpressed NPs in MDCK cells. NP and NPM1 were immuno-stained after the protease treatment at 10 h post-transfection (hpt). Scale bars, 20 µm. Representative images from two independent experiments. **c,** Replication and transcription efficiencies of the reconstituted RNPs, measured by strand-specific RT-qPCR. HEK293T cells were transfected with PB2, PB1, PA, NP proteins, and HA vRNA expression plasmids and the total RNA was extracted at 48 hpt. Their HA vRNA, cRNA, and mRNA copy numbers were compared with those of the RNPs reconstituted with NP wt using one-way ANOVA with Dunnett’s test; \**P*<0.05, \*\**P*<0.01, UD, undetected. Data are presented as mean±S.D. of three independent experiments with two RT-qPCR assays.

**Extended Data Figure 4.**
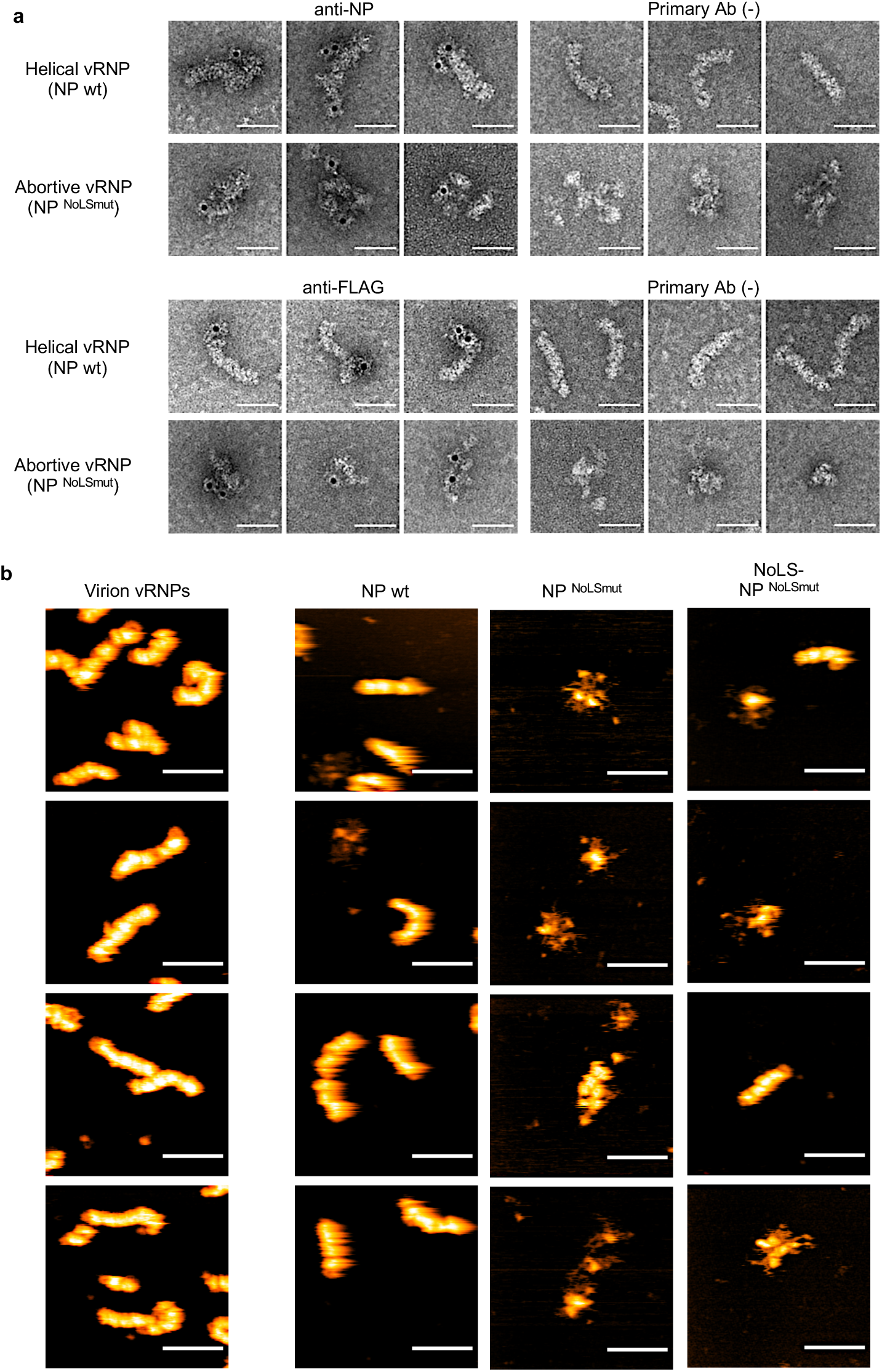
Visualization of the reconstructed vRNP. **a,** Negative-staining immuno-electron microscopy of the purified vRNPs. We analysed each of the 100 labelled vRNPs. The helical vRNPs labelled with anti-NP and anti-FLAG antibodies had one to three gold particles mainly at the terminal region and distributed throughout the vRNPs, respectively. The abortive vRNPs labelled with anti-NP and anti-FLAG antibodies had one to three gold particles. Of 300 or more vRNPs in the primary Ab (-) controls, only one or zero gold particle-bound vRNP was observed. Three representative images are shown. Scale bar, 50 nm. **b,** Supplementary images of the purified vRNPs visualized by HS-AFM. Scale bar, 100 nm.

**Extended Data Figure 5.**
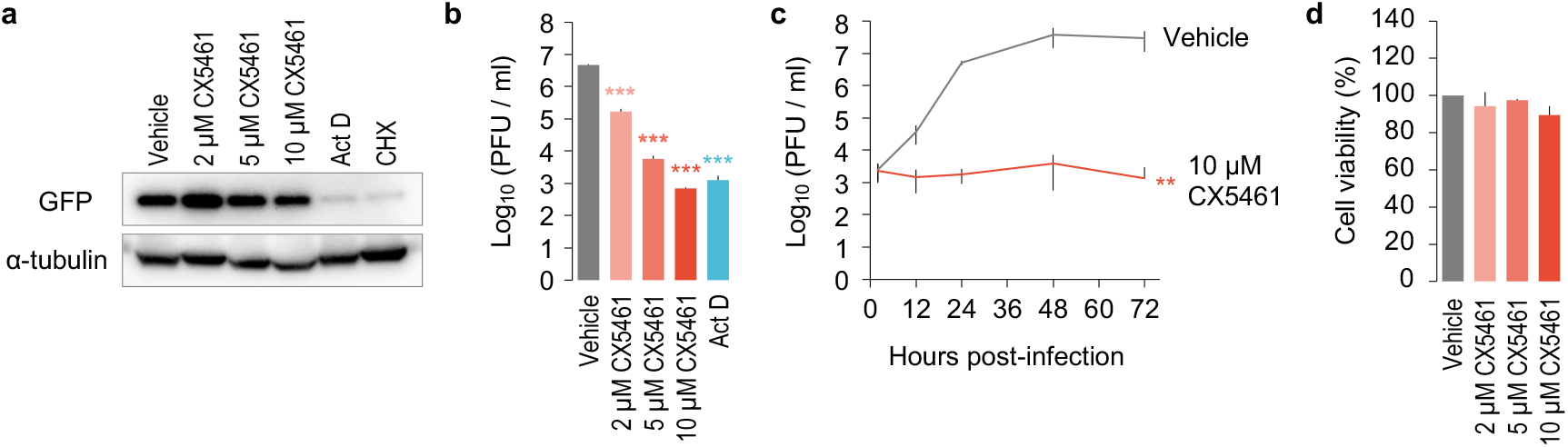
Effect of RNA polymerase I inhibitor on mRNA transcription, viral growth, and cell viability. **a,** Effect on mRNA transcription and translation. A549 cells were transfected with a GFP expression plasmid (pCAGGS/GFP) and incubated in a medium containing CX5461, 10 µg/mL actinomycin D (Act D), 10 µM cycloheximide (CHX), and 1% DMSO (Vehicle) at 12 hpt. After additional 12-h incubation (24 hpt), cell lysate was analysed by western blotting. Representative images from two independent experiments. **b,** Effect on viral growth. A549 cells were pretreated with CX5461 or 10 µg/mL actinomycin D (Act D) for 2 h, followed by virus infection (MOI=0.1). The supernatants were obtained at 24 hpi and subjected to plaque assay. The viral titres were compared with those of vehicle-treated cells using one-way ANOVA with Dunnett’s test; \*\*\**P*<0.001. Data are presented as mean±S.D. of three independent experiments. **c,** Viral growth kinetics in CX5461-treated cells. A549 cells were pretreated with 10 µM CX5461 or vehicle for 2 h, followed by wild-type virus infection (MOI=0.1). The supernatants were obtained at 2, 12, 24, 48, 72 hpi and subjected to plaque assay. The viral titres were compared with those of the vehicle-treated cells using two-way ANOVA; \*\**P*<0.01. Data are presented as mean±S.D. of three independent experiments. **d,** Cytotoxicity of CX5461. A549 cells treated with CX5461 or vehicle for 48 h were subjected to a cell viability assay. The cell viabilities were compared using one-way ANOVA (*P*=0.88). Data are presented as mean±S.D. of three independent experiments.

**Extended Data Figure 6.**
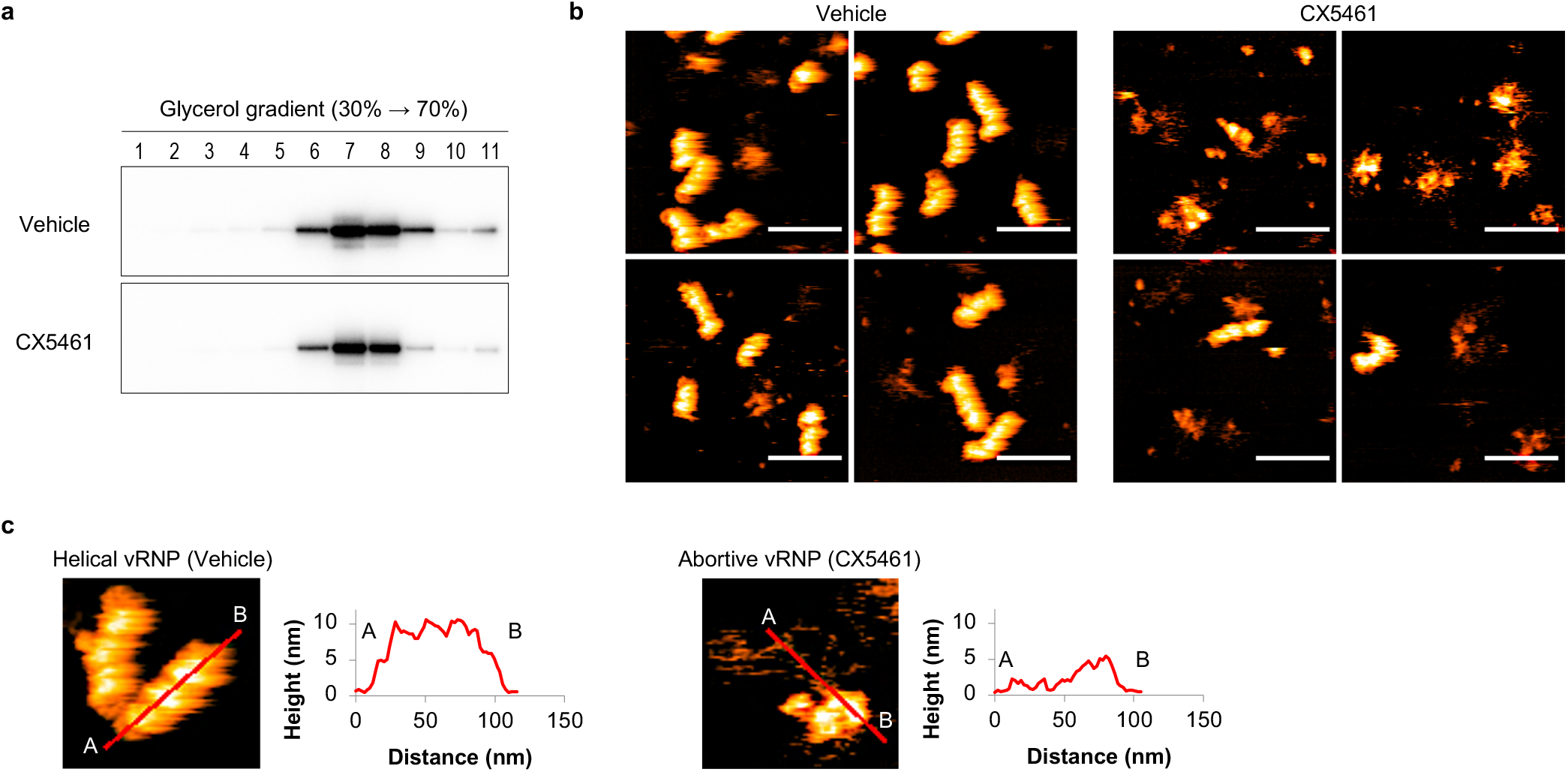
High-speed atomic force microscopy (HS-AFM) analysis of the vRNP purified from the influenza virus-infected cells. **a,** Purification of vRNP. The immunoprecipitated vRNPs from the PB2-FLAG virus-infected cells were further purified by ultracentrifugation through 30% to 70% glycerol gradients. Each fraction was gel-electrophoresed and immunoblotted with anti-NP antibody. **b,** Representative images of the purified vRNPs from the PB2-FLAG virus infected-cells visualized by HS-AFM. Scale bar, 100 nm. **c,** Section analysis of the helical and Abortive vRNPs. Enlarged HS-AFM images of Fig. 4f are shown. Heights of the helical and the abortive vRNPs were measured at the red lines from A to B.

**Extended Data Figure 7.**
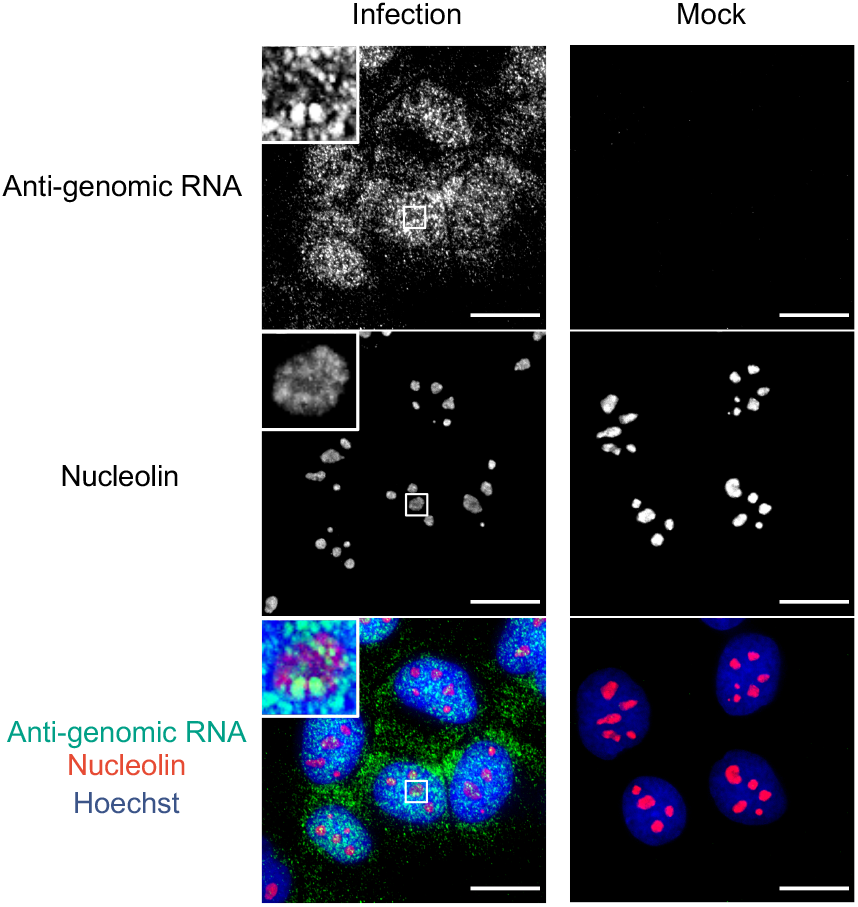
Nucleolar localization of the anti-genomic RNA in influenza virus-infected cells. Anti-genomic RNA containing cRNA and mRNA of the PB2 segment were detected in the virus-infected cells (MOI=5) at 5 hpi by fluorescence *in situ* hybridization. Nucleolin was immuno-stained. Insets: enlarged versions of the selected regions indicated by the white boxes. Scale bars, 20 µm. Representative images from three independent experiments.

## Notes

### Competing Interest Statement

The authors have declared no competing interest.

